# Cardiolipin modulation of MmpL3 in mycobacteria

**DOI:** 10.64898/2026.06.25.734600

**Authors:** Sizhun Li, Chelsea M. Brown, Ruby Hao Sun, Phillip J. Stansfeld, Shu-Sin Chng

## Abstract

The complex cell envelope of mycobacteria is characterized by the presence of a unique outer membrane (OM) rich in mycolic acids (MAs). These long-chain, branched fatty acids confer extreme hydrophobicity to the OM, in part rendering the mycobacterial envelope impermeable to antibiotics and host defences. How MAs are transported from the inner membrane (IM) to the OM is largely unknown. The integral membrane protein MmpL3 plays an essential role in this process, but the mechanism(s) by which it exploits the proton motive force to flip and/or release MAs at the IM, in the form of trehalose monomycolates (TMMs), remains elusive. Here, we reconstitute and quantify the proton translocation activity of MmpL3 in artificial lipid bilayers, and discover a novel role for the phospholipid species, cardiolipin (CL), in regulating MmpL3 function. We find that mutations in conserved residues, or binding of known inhibitors in the central channel of MmpL3 do not diminish proton translocation activity. Instead, the specific presence of CL abolishes proton translocation by MmpL3 in vitro. Furthermore, we establish that an MmpL3 variant containing substitutions in a CL-binding site predicted in silico is no longer modulated by CL in vitro, and is unable to support growth of *Mycobacterium smegmatis*. Our work provides previously unappreciated insights into lipid regulation of MmpL3 activity in mycobacteria, and expands the guiding principles for the development of anti-mycobacterial inhibitors targeting this essential transporter.

**Significance:** Mycobacterial species, including the human pathogen *Mycobacterium tuberculosis*, are surrounded by a double-membrane cell envelope that makes them intrinsically resistant to many antibiotics. Specifically, the outer membrane (OM) contains unique lipids called mycolic acids (MAs), whose transport pathway across the envelope is poorly understood. In this study, we characterized the biochemical activity of the essential MA transporter MmpL3, and uncovered a novel mechanism for lipid-mediated functional regulation. Our work highlights the importance of protein-lipid interactions in defining transporter activity, provides insights into MA transport and OM assembly in mycobacteria, and sets the stage for the development of anti-mycobacterial strategies.

## Introduction

Mycobacteria constitute a group of bacterial pathogens recognized for their global public health impact. *M. tuberculosis*, *M. leprae* and non-tuberculous mycobacteria, such as *M. abscessus*, are responsible for tuberculosis (TB), leprosy and other opportunistic infections in humans, respectively, posing considerable health challenges^1^. Notably, according to the WHO Global Tuberculosis Report, TB accounted for ∼1.6 million deaths out of 10.6 million new cases in 2022 worldwide^2^. Treating TB is challenging, inefficient, and costly; the standard regimen requires daily intake of a combination of four antibiotics – isoniazid (INH), rifampin (RIF), pyrazinamide (PZA), and ethambutol (EMB) – over an extended period, typically six months or longer^1^. Furthermore, the emergence of multi-drug-resistant TB (MDR-TB) and extensively drug-resistant TB (XDR-TB) presents even greater threats to the existing public health system^3^.

The difficulty in combatting mycobacterial infections stems in part from their complex cell envelope structure, which is characterized by its abundance of mycolic acids (MAs)^4^. MAs are exceptionally long fatty acids forming a protective hydrophobic outer membrane (OM) barrier that hinders the effectiveness of many conventional antibiotics. They are synthesized as glycolipids at the inner membrane (IM), and must be transported across an aqueous periplasmic space to be assembled into the OM. MAs are first produced in a form linked to trehalose^5–7^, termed trehalose monomycolate (TMM), via a well-documented biosynthetic pathway^8,9^. These TMMs are then flipped across, and possibly also released from, the IM by the essential membrane protein MmpL3^10^. In addition, TMMs may be acetylated by the putative acetyltransferase TmaT^11^, but how this modification fits within the pathway has remained unclear. While the details of their transport mechanism from the IM to the OM are largely uncharacterized, it is known that the mycolic chain from one TMM molecule is transferred to another TMM molecule to form trehalose dimycolate (TDM), or to the arabinogalactan-peptidoglycan matrix to afford the mycolyl-arabinogalactan-peptidoglycan (mAGP) complex^12^. These acyl transfer reactions are catalyzed by the Ag85 enzymes at the OM.

MmpL3 belongs to the Resistance, Nodulation, and Division (RND) superfamily of membrane transporters^13,14^, and is believed to be powered by the proton motive force (*pmf*)^15^. It has emerged as a significant antibiotic target due to its susceptibility to inhibition by various compounds with different chemical scaffolds. Treatment with direct MmpL3 inhibitors, such as AU1235^16^ or BM212^17^, led to a greatly reduced accumulation of TMMs in the outer leaflet of the IM in mycobacterial spheroplasts, establishing MmpL3 as the TMM flippase^10^. MmpL3 comprises a transmembrane domain with 13 helices, a periplasmic domain, and a cytoplasmic C-terminal domain ^18–20^, though it is unclear whether it functions as a monomer or oligomer^21,22^. MmpL3 interacts specifically with TtfA, whose function is also unknown^23^. Structural characterization revealed that the periplasmic domain contains cavities that bind detergents or lipids, including TMM, suggesting a possible role also in the release of TMM from the IM (post flipping)^24^. Consistent with this idea, a second TMM binding site in the outer leaflet of the transmembrane region of MmpL3 has been identified, and a route by which TMM may move from this site to the periplasmic domain has been proposed^24^. Furthermore, a recent in vitro study demonstrated the possible ability of MmpL3 in releasing phospholipids, albeit of non-native structures, from the bilayer^25^. Presumably, additional factors would be required to shuttle TMM across the periplasmic space.

How MmpL3 utilizes the *pmf* to flip TMM across the IM, and possibly release from the membrane, is not clear. MmpL3 possesses a large central channel demarcated by four transmembrane helices, providing a likely route for proton passage^18^. At the midpoint of this channel, two highly conserved Asp-Tyr pairs (D256-Y646, D645-Y257 in *M. smegmatis* MmpL3), resembling Asp-Asp-Lys and Asp-Asp-Thr triads found in RND transporters AcrB and SecDF^26^, are important for overall function, and believed to mediate proton translocation^22^. Indeed, the channel is a hot spot for binding by putative MmpL3 inhibitors^18^; while this observation suggests proton translocation disruption as a plausible mode of action, it also highlights the possibility that these compounds may impact conformational changes required for TMM transport. The mechanisms of MmpL3 function and inhibition remain to be clarified.

To study MmpL3, we set out to reconstitute its activities in vitro. Here, we establish MmpL3 proton translocation activity in proteoliposomes, and show that this activity is in fact modulated by cardiolipin (CL), a relatively abundant phospholipid species in mycobacteria^27^. Using a quantitative proton translocation assay, we compare purified *M. smegmatis* MmpL3 WT and variants, and find that mutations of the aspartates in the DY pairs do not reduce proton translocation activity. Intriguingly, we show that the presence of CL in proteoliposomes inhibits the passage of protons through MmpL3, indicating a regulatory role. Furthermore, molecular dynamics (MD) simulation-guided mutagenesis of a potential CL-binding site on MmpL3 abolishes CL modulation in proteoliposomes, and compromises its ability to support growth. We demonstrate that the same mutation in *M. tuberculosis* MmpL3 also causes loss of function, suggesting conservation of this CL modulation mechanism. Our work highlights novel insights into the functional regulation of the TMM transporter MmpL3, and opens up new strategies towards the development of anti-mycobacterial inhibitors against this important drug target.

## Results

### MmpL3 facilitates quantifiable proton translocation activity in vitro

To measure proton translocation activity of MmpL3, we expressed and purified C-terminally His_8_-tagged *M. smegmatis* MmpL3 (MmpL3-His) from *Escherichia coli*, and reconstituted the recombinant protein into liposomes composed of synthetic lipids (POPE:POPG = 3:1). MmpL3-His was inserted into proteoliposome membranes in a right-side-out manner, consistent with its orientation in cells, as judged by the inability of carboxypeptidase A (CPA) to access and degrade the C-terminal His_8_ tag (**Figure S1**). The pH-sensitive dye pyranine was encapsulated in the lumen of our proteoliposomes to enable measurement of internal pH (pH_i_) using fluorescence; as protons are translocated across the membrane, the fluorescence emission ratio of pyranine when excited at 450 nm and 400 nm (i.e. I_450_/I_400_) will change according to pH_i_^28^. To drive the movement of protons across MmpL3, we created an artificial pH gradient across the membrane by diluting proteoliposomes (pH_i_ = 7.0) into buffer at pH 6.0. We monitored the rate of lumen acidification in the presence and absence of the potassium (K^+^) ionophore valinomycin, which can prevent the build-up of membrane potential leading to slowing of proton flow (**Figure 1**). Without valinomycin, the I_450_/I_400_ ratio (i.e. pH_i_) stayed essentially constant in empty liposomes after the external pH jump (i.e. no proton leakage) (**Figure 1A**). Even though inserting MmpL3 gave a discernable rate in the change of fluorescence ratio, the effect was very small and hardly quantifiable, similar to what was observed in a previous study^29^. In contrast, while adding valinomycin increased the baseline proton leakage rate in empty liposomes, doing so more significantly enhanced proton translocation in MmpL3 proteoliposomes (by ∼2.6 fold over empty liposomes across 6 biological replicates) (**Figures 1A** and **1B**). We observed similar enhancement in proton translocation when the pH gradient was reversed (i.e. outside pH, or pH_o_ = 8.0) (**Figure S2**). We conclude that MmpL3 demonstrates quantifiable proton translocation activity in vitro, dependent on the direction of the applied pH gradient.

**Figure 1.**
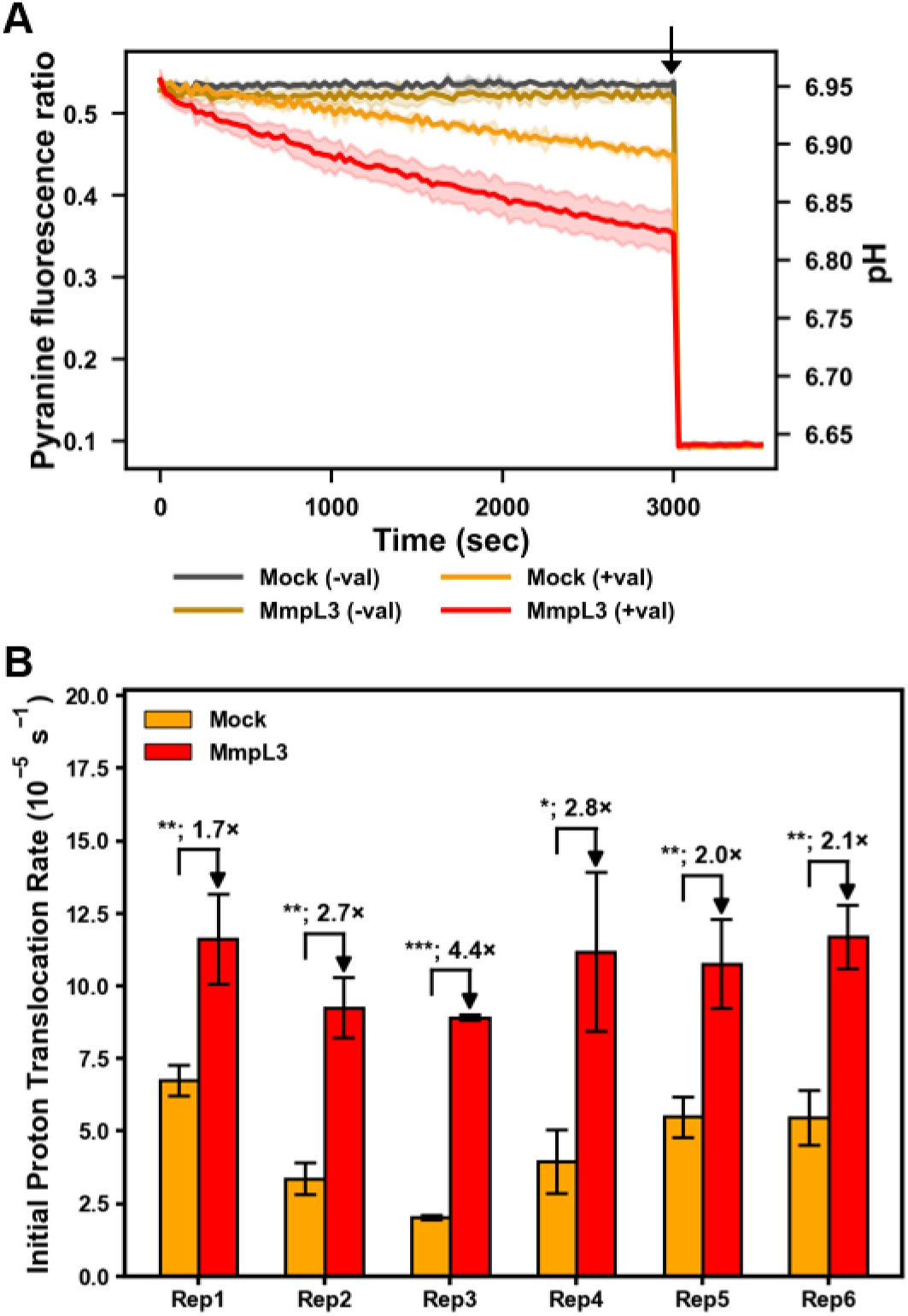
Purified MmpL3 exhibits proton translocation activity in proteoliposomes comprising synthetic lipids (POPE:POPG = 3:1). (A) Representative proton translocation assays showing change of pyranine fluorescence ratios (I_450_/I_400_) over time of empty (mock) and MmpL3-embedded liposomes due to lumenal acidification in the presence of an artificial pH gradient (initial pH_i_ = 7.0; pH_o_ = 6.0). Corresponding pH_i_ values (using a calibration curve) are shown on the right axis. MmpL3-His (0.6 pmol) was reconstituted into liposomes composed of synthetic lipids (POPE:POPG = 3:1). Assays were performed in the presence or absence of 1 µM valinomycin as indicated. 1 µM nigericin was added (*black arrow*) to disrupt the proton gradient. Means and standard deviations (±SD) of fluorescence ratios from measurements of technical repeats are plotted. (B) Initial mean “proton translocation” rates in empty (mock) and MmpL3-embedded liposomes were quantified (from technical repeats such as in (A)) and displayed for six biological replicates (i.e. independently reconstituted proteoliposomes). For each biological replicate, initial rates were calculated by analyzing the initial linear slopes of fluorescence ratio curves, averaged across 2-3 technical repeats. Error bars represent ±SD of initial slopes. Fold change with or without MmpL3 within each replicate is displayed. Student’s t-test: *, p < 0.05; **, p < 0.01; ***, p < 0.005.

Taking advantage of our in vitro assay, we went ahead to test whether mutations in the highly conserved Asp-Tyr (DY) pairs in MmpL3^30^, which are believed to be essential for proton translocation, impacted activity. We first showed that D256A and D645A mutants were unable to rescue the growth of an *mmpL3* conditional knock-out (cKO) strain depleted of MmpL3 (**Figure S3**), indicating these mutations abolished function. In contrast, Y257F and Y646F mutants effectively restored growth. Unexpectedly, we found that all four single variants exhibited substantial proton translocation activities when purified and reconstituted into proteoliposomes (**Figure 2**). Specifically, Y257F displayed similar proton translocation levels to WT MmpL3, while D256A, D645A and Y646F mutants exhibited even higher activities, which did not correlate with their observed (non)-functionality in cells. These findings suggested that the DY pairs in MmpL3 could play additional roles beyond simply translocating protons across the IM.

**Figure 2.**
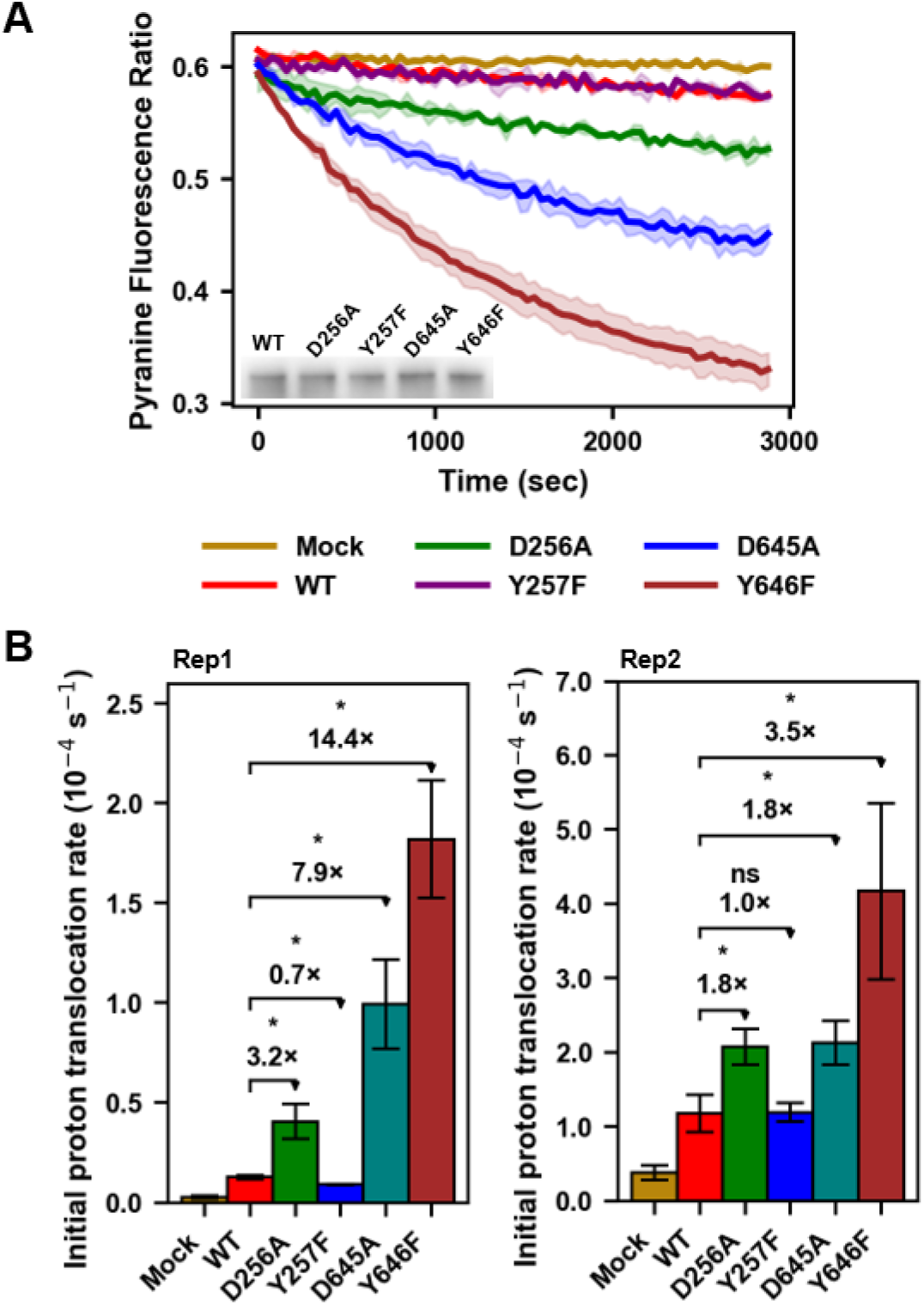
Purified MmpL3 channel DY-pair variants display comparable or higher proton translocation activity in proteoliposomes compared to wild-type MmpL3. (A) Representative proton translocation assays showing change of pyranine fluorescence ratios (I_450_/I_400_) over time of empty (mock) liposomes and proteoliposomes reconstituted with indicated MmpL3 channel variants, due to lumenal acidification in the presence of an artificial pH gradient (initial pH_i_ = 7.0; pH_o_ = 6.0). Purified MmpL3-His variants (0.6 pmol) were reconstituted into liposomes composed of synthetic lipids (POPE:POPG = 3:1). Inset shows α-His immunoblot analysis of reconstituted MmpL3 proteoliposomes used. Assays were performed in the presence of 1 µM valinomycin. Means and standard deviations (±SD) of fluorescence ratios from measurements of technical repeats are plotted. (B) Initial mean “proton translocation” rates in empty (mock) liposomes and proteoliposomes reconstituted with indicated MmpL3 variants were quantified (from technical repeats such as in (A)) and displayed for 2 of 3 biological replicates (i.e. independently reconstituted proteoliposomes) performed. For each biological replicate, initial rates were calculated by analyzing the initial linear slopes of fluorescence ratio curves, averaged across 2-3 technical repeats. Error bars represent ±SD of initial slopes. Fold changes comparing each MmpL3 DY-pair variant to MmpL3^WT^ within each replicate are displayed. Student’s t-test: *, p < 0.05; ns, not significant.

We also examined the effects of known MmpL3 inhibitors, AU1235^16^, BM212^17^ and NITD-304^31^, since these compounds likely bind directly in the central channel and are believed to inhibit proton translocation. To our surprise, the proton translocation activity of MmpL3 was essentially unaffected in the presence of BM212, AU1235, or NITD304 (**Figure 3**). For the latter two, we employed concentrations estimated to drive at least ∼25% of reconstituted MmpL3 into inhibitor-bound forms, when considering 3D diffusion (**Table S1**); due to the propensity of these inhibitors to partition into the lipid bilayer, fraction of bound-MmpL3 was expected to be much higher. Beyond these concentrations, these compounds began to affect proton leakage in empty liposomes non-specifically, precluding useful interpretation. These observations challenge the initial assumption that channel binding molecules must impact proton translocation^18^. Since AU1235 and BM212 compounds have been demonstrated to inhibit the TMM flipping activity of MmpL3 in *M. smegmatis* spheroplasts^10^, we propose that they might instead affect how proton translocation is coupled to lipid transport across the membrane.

**Figure 3.**
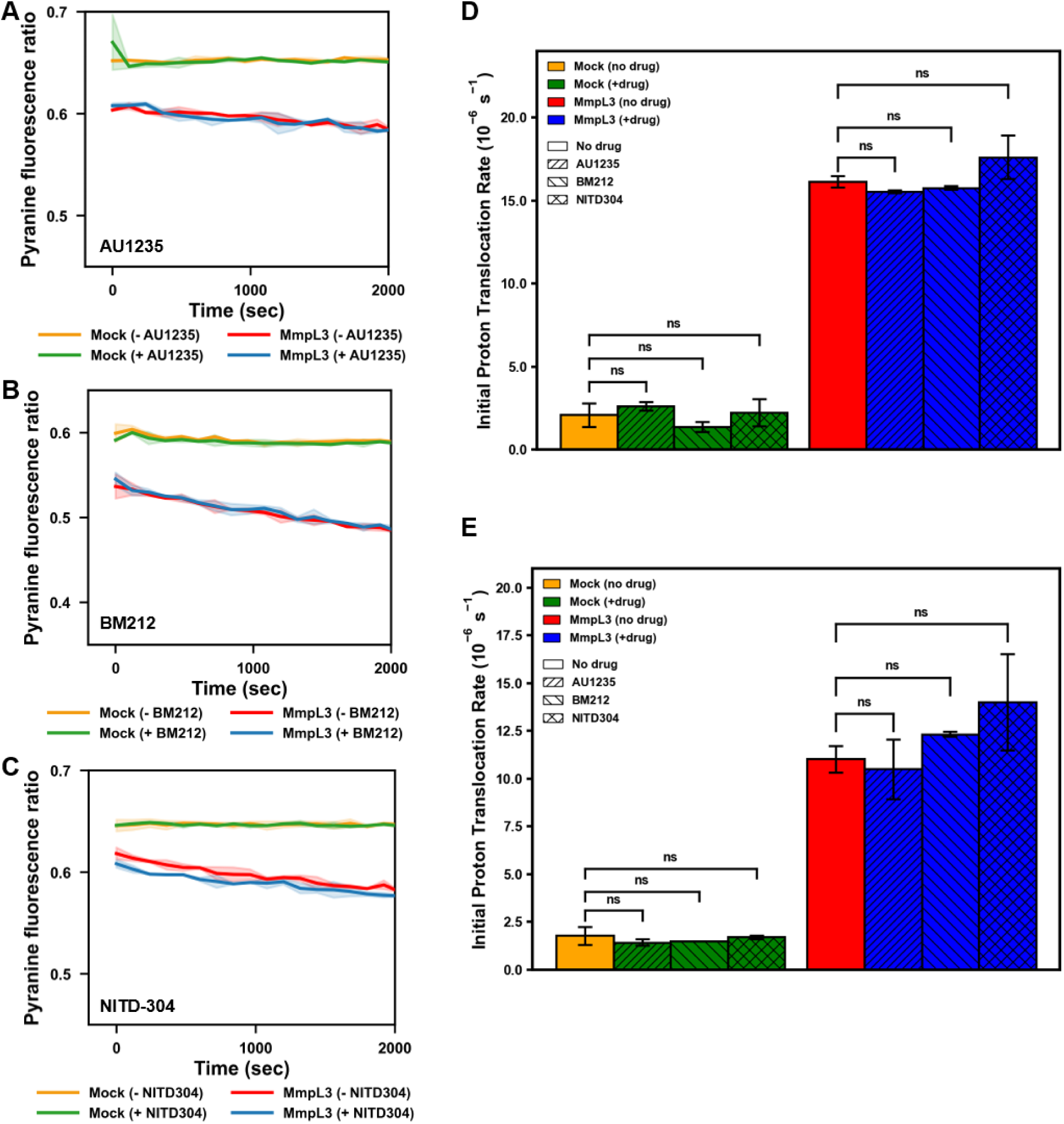
Known chemical inhibitors do not reduce proton translocation activity of purified MmpL3 in proteoliposomes. (A-C) Representative proton translocation assays showing the change of pyranine fluorescence ratios (I_450_/I_400_) over time of empty (mock) and MmpL3-embedded liposomes in the presence or absence of (A) AU1235 (0.1 µM), (B) BM212 (1.0 µM), or (C) NITD-304 (0.05 µM), due to lumenal acidification in the presence of an artificial pH gradient (initial pH_i_ = 7.0; pH_o_ = 6.0). MmpL3-His (0.6 pmol) was reconstituted into liposomes composed of synthetic lipids (POPE:POPG = 3:1). Assays were performed in the presence of 1 µM valinomycin. Means and standard deviations (±SD) of fluorescence ratios from measurements of technical repeats are plotted. (D-E) Initial mean “proton translocation” rates in empty (mock) and MmpL3-embedded liposomes, in the presence or absence of indicated inhibitor, were quantified (from technical repeats such as in (A-C)) and displayed for 2 of 3 biological replicates (i.e. independently reconstituted proteoliposomes) performed. For each biological replicate, initial rates were calculated by analyzing the initial linear slopes of fluorescence ratio curves, averaged across 2 technical repeats. Error bars represent ±SD of initial slopes. Student’s t-test: ns, not significant. At the concentrations used, AU1235, BM212 and NITD-304 did not significantly impact proton translocation rates in empty (mock) or MmpL3-embedded liposomes, despite AU1235 and NITD-304 estimated to occupy >25% MmpL3 binding sites (see **Table S1**).

### MmpL3 proton translocation activity is modulated by cardiolipin

The lipid environment can significantly impact membrane protein activity. To mimic a more native lipid environment for MmpL3 and its mutants, we also investigated proton translocation activity in proteoliposomes containing extracted lipids from *M. smegmatis* cells (80% POPE:POPG (3:1), 20% w/w *M. smegmatis* polar lipids) (**Figures 4A** and **4B**). Incorporation of *M. smegmatis* lipids significantly increased baseline proton leakage in empty liposomes. Curiously, however, we no longer detected additional proton translocation activity in WT MmpL3 proteoliposomes containing these *M. smegmatis* lipids, indicating possible modulation by specific lipid species. MmpL3 has been shown to co-purify with phosphatidylethanolamine (PE), phosphatidylglycerol (PG), phosphatidylinositol (PI), diacylglycerol (DAG) and cardiolipin (CL)^32^. To test whether any of these lipids can modulate MmpL3, we went on to measure its proton translocation activity in proteoliposomes containing PI, DAG or CL at physiologically relevant concentrations^27^. 10% w/w PI or 2% w/w DAG did not significantly affect MmpL3’s proton translocation activity (**Figures 4C**, **4D**, **4F** and **4G**). In contrast, 20% w/w CL was found to strongly modulate MmpL3 activity, completely inhibiting its proton translocation (**Figures 4E** and **4H**). Consistent with this observation, liposomes with 100% *E. coli* polar lipids (PE:PG:CL = ∼67:23:10) also showed no detectable proton translocation by MmpL3 (**Figures 4I** and **4J**). We conclude that CL may modulate MmpL3 function. Our findings raised the question of whether the MmpL3^D256A^ or MmpL3^D645A^ variants were non-functional due to changes in CL modulation; yet we found that CL continued to exert an inhibitory effect on the proton translocation activities of either mutant in proteoliposomes (**Figure S4**), again implying complex influence on MmpL3 function by the DY pairs.

**Figure 4.**
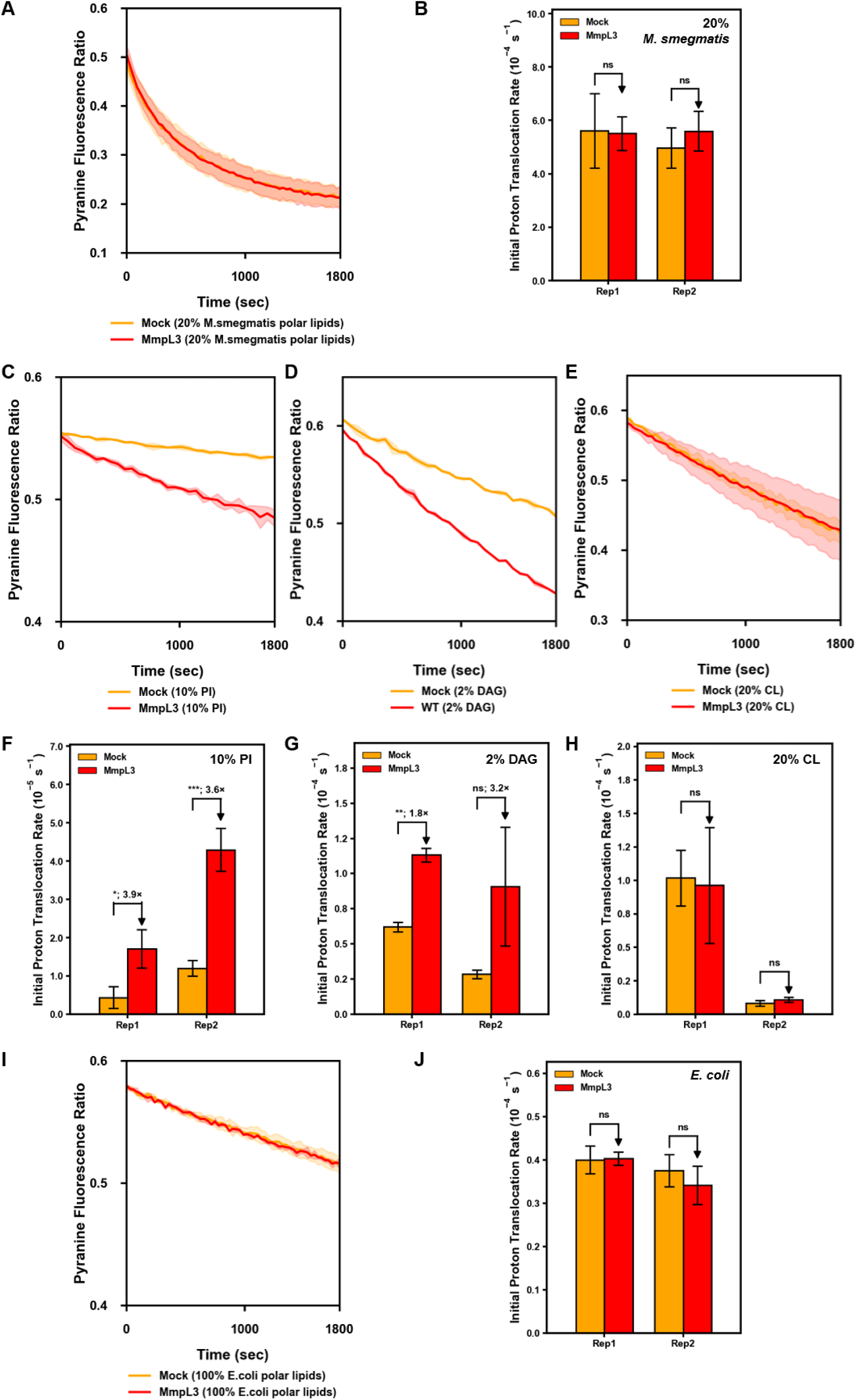
The presence of cardiolipin inhibits proton translocation activity of MmpL3 in proteoliposomes. (A, C-E, I) Representative proton translocation assays showing the change of pyranine fluorescence ratios (I_450_/I_400_) over time of empty (mock) and MmpL3-embedded liposomes due to lumenal acidification in the presence of an artificial pH gradient (initial pH_i_ = 7.0; pH_o_ = 6.0). MmpL3-His (0.6 pmol) was reconstituted into liposomes comprising (A) 20% w/w *M. smegmatis* lipids, (C) 10% w/w PI, (D) 2% w/w DAG, or (E) 20% w/w CL in POPE:POPG (3:1), or (I) 100% *E. coli* polar lipids. Assays were performed in the presence of 1 µM valinomycin. Means and standard deviations (±SD) of fluorescence ratios from measurements of technical repeats are plotted. (B, F-H, J) Initial mean “proton translocation” rates in empty (mock) and MmpL3-embedded liposomes were quantified (from technical repeats such as in (A, C-E, I)) and displayed for 2 of 3 biological replicates (i.e. independently reconstituted proteoliposomes) performed. For each biological replicate, initial rates were calculated by analyzing the initial linear slopes of fluorescence ratio curves, averaged across 2-4 technical repeats. Error bars represent ±SD of initial slopes. Fold change with or without MmpL3 within each replicate is displayed. Student’s t-test: *, p < 0.05; **, p < 0.01; ***, p < 0.005; ns, not significant.

### Mutation of a potential binding site on MmpL3 abolishes modulation by cardiolipin

To understand the mechanism(s) by which CL modulates proton translocation activity, we attempted to identify potential CL binding sites on MmpL3. We applied coarse-grained molecular dynamics (CGMD) simulations to *M. smegmatis* MmpL3 in a bilayer comprising PE, PG, and CL (at 75:20:5 ratio). Over 50 µs of simulation time, we found four distinct CL binding sites, each containing multiple residues (**Figure S5A and S5B**); these sites exhibited high CL occupancy (50-62%) throughout the simulations, but saw significant reductions when selected residues were substituted with alanines in silico (**Figure S5B**).

To test the functional significance of these potential CL-binding sites, we expressed MmpL3 alanine-substituted variants for each site, and tested their abilities to support the growth of the *mmpL3* cKO strain. Unexpectedly, all these variants still effectively complemented growth, implying that alanine substitutions did not adversely affect MmpL3 function (**Figure S5C**). To introduce more drastic changes at these sites, we then engineered new variants where lysine/arginine residues within each site were mutated to glutamates. Intriguingly, glutamate substitutions of R392, K399 and R400 on Site 4 (S4EEE), which are highly conserved, resulted in a loss of growth complementation of the cKO strain (**Figures 5A** and **5B**), indicating that these three residues are functionally important, possibly in CL binding. Moreover, MmpL3^S4EEE^ displayed a dominant negative effect even though it was expressed at lower levels (**Figure S5D**), evident from growth inhibition even when WT MmpL3 was present (**Figure 5B**). While individual or double combinations of R392E, K399E and R400E did not render *M. smegmatis* MmpL3 non-functional (**Figures S5C** and **S5D**), we further demonstrated that expressing *M. tuberculosis* MmpL3^K394E/R395E^ and MmpL3^S4EEE^ in the *M. smegmatis mmpL3* cKO strain also prevented growth (**Figure S6**), suggesting that the function of these specific residues is likely conserved in mycobacterial MmpL3.

**Figure 5.**
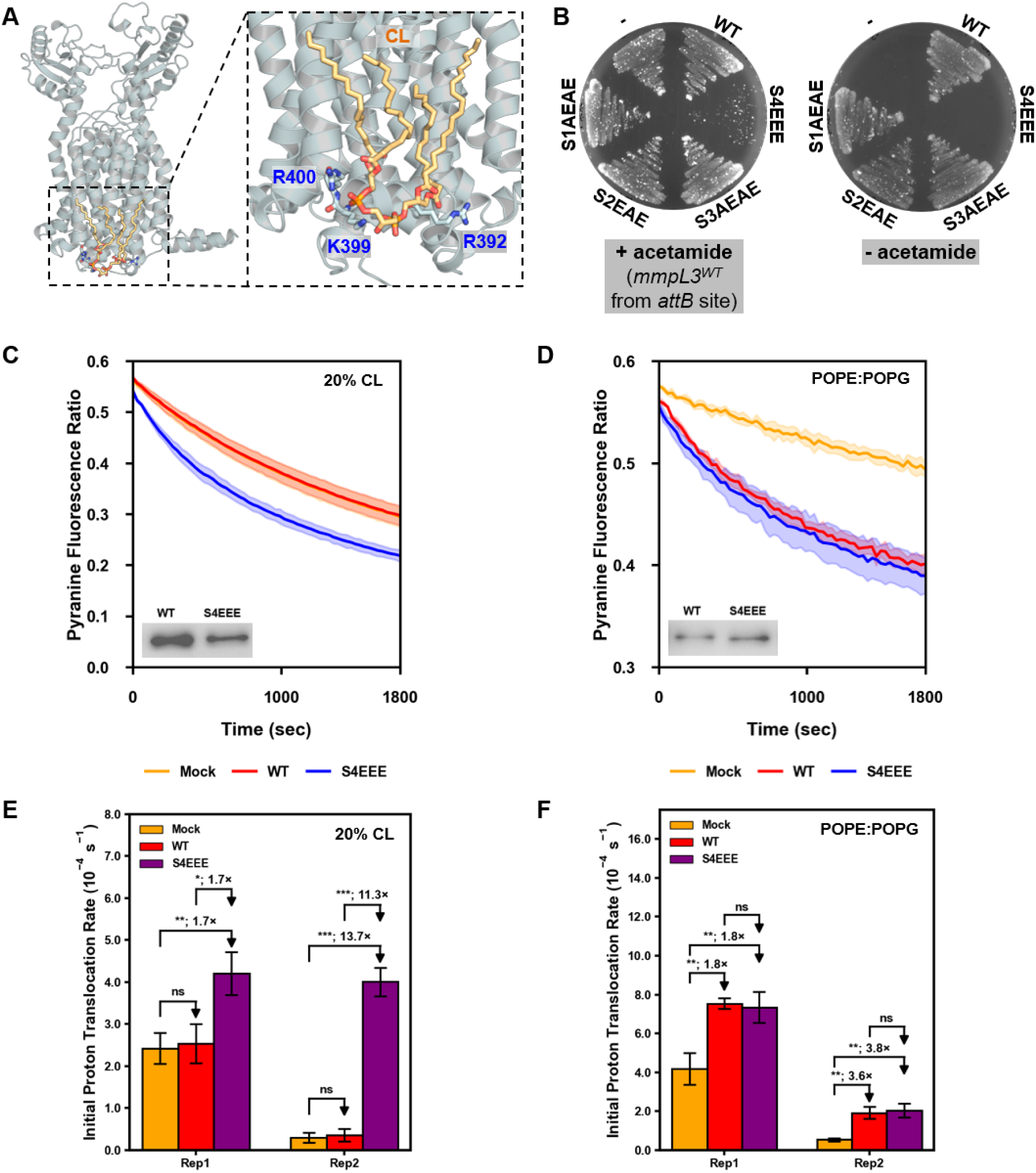
The proton translocation activity of a non-functional MmpL3^S4EEE^ variant with charge-swap substitutions in a putative cardiolipin binding site is no longer inhibited by the presence of cardiolipin. (A) Cartoon representation of MmpL3 (PDB 7N6B) illustrating the putative cardiolipin binding site (site 4) comprising R392, K399 and R400. (B) Growth complementation by *M. smegmatis* MmpL3^WT^ or MmpL3 variants (with substitutions in putative CL binding sites) expressed *in trans* from pMV261 in the *mmpL3* conditional knockout (cKO) strain. Expression of *mmpL3^WT^* from the *attB* site was induced by addition of acetamide (0.2% w/v) where indicated. MmpL3 variants tested: S1AEAE – T236A/K313E/Y317A/K563E; S2EAE – K375E/T376A/R677E; S3AEAE – W737A/R740E/W741A/R744E; S4EEE – R392E/K399E/R400E. (C-D) Representative proton translocation assays showing change of pyranine fluorescence ratios (I_450_/I_400_) over time of empty (mock) liposomes and proteoliposomes reconstituted with MmpL3^WT^ or MmpL3^S4EEE^ variant, due to lumenal acidification in the presence of an artificial pH gradient (initial pH_i_ = 7.0; pH_o_ = 6.0). Purified MmpL3 variants (0.6 pmol) were reconstituted into liposomes comprising (C) 20% w/w CL in POPE:POPG (3:1), or (D) 100% POPE:POPG (3:1). Insets show α-His immunoblot analyses of reconstituted MmpL3 proteoliposomes used. Assays were performed in the presence of 1 µM valinomycin. Means and standard deviations (±SD) of fluorescence ratios from measurements of technical repeats are plotted. (E-F) Initial mean “proton translocation” rates in empty (mock) liposomes and proteoliposomes reconstituted with indicated MmpL3 variants were quantified (from technical repeats such as in (C-D)) and displayed for 2 of 3 biological replicates (i.e. independently reconstituted proteoliposomes) performed. For each biological replicate, initial rates were calculated by analyzing the initial linear slopes of fluorescence ratio curves, averaged across 2-3 technical repeats. Error bars represent ±SD of initial slopes. Fold changes comparing MmpL3^S4EEE^ to MmpL3^WT^ or mock within each replicate are displayed. Student’s t-test: *, p < 0.05; **, p < 0.01; ***, p < 0.005; ns, not significant.

We hypothesize that the S4EEE mutations cause loss of CL binding and modulation on proton translocation, leading to dysregulation of MmpL3 and thus a dominant negative effect. In CGMD simulations, S4EEE variants also had significantly reduced interactions with CL for both *M. smegmatis* and *M. tuberculosis* MmpL3 (**Figure S5B**). We therefore expressed and purified *M. smegmatis* MmpL3^S4EEE^ from *E. coli* cells and studied its proton translocation activity in proteoliposomes with or without CL. Remarkably, we found that while MmpL3^S4EEE^ activity without CL was similar to that of MmpL3^WT^, it remained unaffected in the presence of CL (**Figures 5C-F**). This observation is consistent with the loss of CL binding, hence modulation. We reasoned that this dysregulation causes the observed dominant negative effect when MmpL3^S4EEE^ was expressed even in WT cells. Indeed, expressing MmpL3^S4EEE^, but not MmpL3^WT^, in *M. smegmatis* cells disrupted the pH gradient across the IM (**Table 1**), likely inhibiting growth. In conclusion, our findings suggest that CL binds MmpL3 at the site comprising R392, K399, and R400, exerting a role in modulating proton translocation activity.

**Table 1.** MmpL3^S4EEE^ disrupted the pH gradient across the IM in *M. smegmatis* cells.

| MmpL3 | Induction | Drug<sup>a</sup> | pH outside | pH inside | ΔpH |
| --- | --- | --- | --- | --- | --- |
| S4EEE | - | DMSO | 6.8 | 6.96 ($\pm 0.02$) | 0.16 |
| S4EEE | + | DMSO | 6.8 | 6.83 ($\pm 0.00$) | 0.03 |
| S4EEE | - | CCCP<sup>a</sup> | 6.8 | 6.80 ($\pm 0.01$) | 0.00 |
| S4EEE | + | CCCP | 6.8 | 6.79 ($\pm 0.00$) | -0.01 |
| WT | - | DMSO | 6.8 | 6.99 ($\pm 0.01$) | 0.19 |
| WT | + | DMSO | 6.8 | 6.97 ($\pm 0.01$) | 0.17 |

<sup>a</sup>CCCP (added in DMSO) disrupts the *pmf*, and was used as a control in this experiment.

## Discussion

In this study, we have successfully developed a quantitative assay to demonstrate proton translocation activity of purified MmpL3 in proteoliposomes. Using this assay, we have found that mutations in conserved DY residues, or binding of certain inhibitors, in the central channel of MmpL3 do not in fact reduce proton translocation activity, contrary to expectations. In addition, we have established that MmpL3 proton translocation activity is modulated by CL, possibly through direct protein-lipid interactions at a conserved site in the cytoplasmic leaflet of the IM. Our work revealed a novel regulatory mechanism of MmpL3, which may be exploited for the development of new anti-mycobacterial compounds.

How MmpL3 translocates protons and couples the *pmf* to TMM transport remains unclear. The conserved DY pairs in MmpL3 likely play a role in creating a proton relay pathway^18^, yet key mutations D256A or D645A did not reduce proton translocation in vitro despite being unable to support growth in cells (**Figure 2**). In the case of SecDF from *Thermus thermophilus*, mutation of either of the aspartates in its Asp-Asp-Thr triad greatly reduced proton translocation activity in *E. coli* spheroplasts^26^, more directly defining the requirement of these residues in proton movement across the membrane. Our observations may suggest functional redundancy of the two aspartate residues in MmpL3 in proton translocation, or the presence of alternative proton relay residues that can compensate for loss of either aspartate. Ultimately, however, these aspartate variants do not exhibit transport function in cells; beyond proton translocation, the DY pairs are perhaps more important for coupling the *pmf* to conformational changes required for TMM transport, either flipping and/or release, by MmpL3.

The discovery that CL modulates MmpL3 proton translocation activity is intriguing. In eukaryotic organelles, such as the mitochondria, CL is critical for the functions of respiratory chain complexes, including the F_1_F_0_ ATP synthase^33^. In bacteria, CL is also known to exhibit interactions with a plethora of membrane proteins, including AmtB^34^, SecYEG^35^, and LeuT^36^, possibly promoting function via stabilization of oligomeric states. Interestingly, it is possible for CL binding to directly regulate channel activity in some transporters, such as AqpZ^37^ and KcsA^38^; however, the mechanisms remain poorly understood. We have shown that CL modulates the proton translocation activity of MmpL3 in vitro, potentially via binding at conserved residues R392/K399/R400. These residues are distal from the central channel of MmpL3, implying that CL regulation may occur through binding-induced conformational changes, in a manner that still modulates MmpL3^D256A^ and MmpL3^D645A^ variants exhibiting higher proton translocation activity (**Figure S4**). Of note, recent trimeric structures of another MmpL family protein (MmpL5) revealed lipid species, including CL, bound at/near residues corresponding to the CL-binding site in MmpL3^39,40^, underscoring the possible importance of CL in regulating channel activity and/or oligomerization. While such structural modulation needs to be further characterized, it is clear that the cell has evolved a unique mechanism to tightly control the MmpL3 channel. CL-mediated inhibition of proton translocation may be critical to prevent indiscriminate proton passage and dissipation of the *pmf*, and/or to ensure efficient TMM transport. Consistently, both MmpL3 and anionic lipids (including CL) localize at the poles and division septa in growing cells^41^, perhaps enabling spatially organized interaction and regulation. A recent study has revealed connections between CL and the transport of LPS, the major glycolipid in Gram-negative bacterial OMs^42^. Our work now uncovers an analogous link between CL and TMM/MA transport, highlighting previously unappreciated crosstalk between these major lipid pathways in OM biogenesis.

Understanding the molecular mechanism(s) of MmpL3 inhibitors is pivotal for the development of novel anti-TB drugs. A recent study reported that a TMM analog could stimulate the proton translocation activity of MmpL3 in vitro, and several indole-2-carboxamide compounds inhibited this enhanced activity^29^. However, it was unclear if these compounds interfered directly with proton translocation, or the ability of the TMM analog in binding to, thus stimulating, MmpL3. In our studies, we evaluated several other promising inhibitors, including NITD-304^31^, AU1235^16^, and BM212^17^. Contrary to expectations, none of these compounds demonstrated any inhibitory effect on MmpL3, suggesting that compounds binding in the central channel of MmpL3 might not block or reduce proton translocation. While our results do not provide a definitive explanation for their mechanisms, it is plausible that these inhibitors act on downstream processes of proton translocation, potentially influencing the coupling to TMM flipping and/or its release. The precise mode(s) of action of these MmpL3 inhibitors require further exploration.

## Materials and Methods

### Strains and growth conditions

*Mycobacterium smegmatis* mc^2^155 strains were cultured at 37 °C in Middlebrook 7H9 liquid medium supplemented with 10% ADC enrichment and 0.05% Tween 80, or in LB medium containing 0.05% Tween 80. *M. smegmatis* colonies were grown on 7H10 solid agar supplemented with 10% OADC enrichment, or LB solid agar. Antibiotics used included hygromycin B (50 μg/mL) and kanamycin (25 μg/mL). *E. coli* NovaBlue or DH5α competent cells were used for cloning and *E. coli* BL21(λDE3) cells served as the host for expression of WT MmpL3-His and its variants (see below). *E. coli* cultures were grown in LB medium/agar.

### Generation of *M. smegmatis* Δ*mmpL3 attB*::*p_ace_-mmpL3*, the *mmpL3* conditional knockout strain

The *M. smegmatis mmpL3* conditional knockout strain (Δ*mmpL3 attB*::*p_ace_-mmpL3*) was engineered from the Δ*mmpL3 attB*::*kanR-MOP-mmpL3_Msm_* strain (VM9), previously constructed within our laboratory^43^. This latter strain lacks the chromosomal copy of *mmpL3* but maintains viability through *mmpL3* expressed from an integrative plasmid (pJEB402^44^) at the attB site, using the mycobacterial optimized promoter (*MOP* promoter). To replace the MOP promoter with an acetamide-inducible promoter^45^ (*p_ace_*), we generated a pJEB402 plasmid containing the *mmpL3_Msm_* gene under an acetamide-inducible promoter (pJEB402-*hygR*-*p_ace_*-*mmpL3*, **Table S2**). This construct was introduced via electroporation into the VM9 and transformants were grown on LB plates containing hygromycin. The successful replacement of the *MOP-mmpL3_msm_* allele with *p_ace_-mmpL3_msm_* was confirmed by PCR and subsequent Sanger sequencing. The resulting Δ*mmpL3 attB*::*p_ace_-mmpL3* only grew in the presence of acetamide.

### MmpL3 mutant complementation

To evaluate the functionality of MmpL3 variants, wild-type *mmpL3* was cloned into the pMV261 expression vector (kanamycin resistance), placing the gene under the control of the constitutive *hsp60* promoter (**Table S2**). Mutant alleles were introduced via site-directed mutagenesis (see **Table S3** for primers used). The resulting constructs were introduced into the Δ*mmpL3 attB*::*p_ace_-mmpL3* strain by electroporation, and transformants were selected on LB agar plates containing kanamycin and 0.2% (w/v) acetamide. Individual colonies were subsequently streaked onto LB agar plates containing kanamycin either with or without 0.2% acetamide. In this system, expression of the chromosomally integrated *mmpL3* is induced by acetamide; thus, in the absence of inducer, cell viability depends solely on the plasmid-encoded MmpL3 variant. Functional mutants complemented the conditional knockout and supported growth under both inducing and non-inducing conditions, whereas non-functional mutants supported growth only in the presence of acetamide.

### Overexpression and purification of MmpL3-His

The overexpression and purification of *M. smegmatis* MmpL3-His were conducted with slight modifications to a previously described protocol^10^. Briefly, *E. coli* BL21(λDE3) cells containing pET22/42-*mmpL3-His* were cultured at a scale of 10–20 L and induced at an OD 600 of 0.6–0.8 with 0.5 mM IPTG for 3 h. The harvested cells were resuspended in 200 mL of prechilled buffer A (50 mM potassium phosphate, 150 mM KCl, 5 mM MgCl_2_, pH 8.0) supplemented with 100 μg/mL lysozyme, 100 μM phenylmethylsulfonyl fluoride (PMSF), 1 μM phosphoramidon, and 50 μg/mL DNase I, and subsequently lysed by tip sonication. Membrane fragments were pelleted by ultracentrifugation, and then resuspended in buffer B (50 mM potassium phosphate, 150 mM KCl, 5 mM MgCl_2_, 10% glycerol, 1% DDM, pH 8.0) by gentle shaking on ice for 3 h. The suspension was subsequently clarified by ultracentrifugation at 36,000 × *g* for 1 h, and the supernatant was collected. Subsequent purification was performed using TALON cobalt affinity chromatography. The clarified supernatant was loaded onto 1 mL of pre-equilibrated TALON cobalt resin and incubated on ice for 1 h with gentle mixing. The resin was washed five times with 20 mL of equilibration buffer per wash. Bound protein was eluted with 5 mL of elution buffer (50 mM potassium phosphate, 150 mM KCl, 5 mM MgCl_2_, 10% glycerol, 0.05% DDM, 300 mM imidazole, pH 8.0), and the elution fraction was collected and pooled. Following affinity purification, size-exclusion chromatography was performed in buffer C (50 mM potassium phosphate, 150 mM KCl, 5 mM MgCl₂, 10% glycerol, 0.05% DDM, pH 8.0). Purified MmpL3-His concentrations were quantified using the DC protein assay, with final yields of 0.8–1.5 mg/mL in approximately 1 mL of concentrated protein.

### MmpL3 reconstitution into proteoliposomes

To prepare liposomes, 10 mg of lipids dissolved in chloroform were dried overnight in a fume hood. The dried lipids were hydrated in 1.0 mL of an external buffer solution (composed of 100 mM potassium phosphate, 0.024 M K_2_SO_4_, pH 7.0) containing 0.5 μM pyranine. Lipids were hydrated by repeated freeze–thaw cycles with intermittent bath sonication to promote uniform dispersion. At least 25 freeze–thaw cycles were typically required, and hydration was considered complete when the suspension became optically clear. The dispersed lipids were then extruded through a 200 nm-pore membrane using an Avanti Mini Extruder to produce liposomes of homogeneous size.

To generate MmpL3 proteoliposomes, 1.0 mL of preformed liposomes was incubated with 25.5 µL of 20% (w/v) DDM to a final detergent concentration of 0.5% at 4 °C for 1 h. Subsequently, 10 μg of purified MmpL3-His was added to the liposomes and incubated at 4°C for 2 hours. The sample was subjected to two rounds of incubation with SM-2 BioBeads (BioRad) at 4°C for 1 hour each to remove DDM and facilitate the integration of MmpL3 into the lipid bilayer of the liposomes. Following reconstitution, MmpL3 proteoliposomes were applied to a PD-10 gel filtration column (Cytiva Life Sciences) to eliminate free pyranine and other low-molecular-weight contaminants. Fractions of 200 µL were collected and analyzed by fluorescence, and the three fractions corresponding to the fluorescence peak were pooled (total 600 µL) and used for subsequent assays.

### Proton translocation activity assays

4 μL of MmpL3 proteoliposomes (∼0.067 μg or ∼0.6 pmol MmpL3-His) were loaded into 96-well plates, followed by the addition of 200 μL of an external buffer solution (100 mM potassium phosphate, 24 mM K_2_SO_4_, 1.0 μM valinomycin) at desired pH (pH 6, or pH 8) using a multichannel pipette. The pyranine fluorescence was immediately measured using a plate reader (Molecular Devices, SpectraMax M5), and the fluorescence ratio of I_450_/I_400_ was automatically calculated by the plate reader. To assess the impact of MmpL3 inhibitors, the compounds were pre-mixed with the external buffer solution before initiating the pH jump experiment. The maximum concentrations of inhibitors that did not affect the proton leakage rate of mock proteoliposomes were used – AU1235 (0.1 µM), BM212 (1.0 µM), NITD-304 (0.05 µM). At these concentrations, we estimated >25% occupancy of the available MmpL3-His in proteoliposomes (see **Table S1**). The proton translocation rate was determined by the value of the initial slope of I_450_/I_400_ curve over time, averaged over 2-4 technical repeats in each biological replicate. For each experiment, at least 3 biological replicates (i.e. independently reconstituted proteoliposomes) were performed.

### Molecular dynamics simulations

The structure of *M. smegmatis* MmpL3 was generated based upon the solved experimental structure (PDB ID: 7N6B)^19^. Residues at the turn of the amphipathic helix (residues 359-363) were not resolved in this structure, so the SwissModel online server^46^ was used to complete this section of the structure. The structure of *M. tuberculosis* MmpL3 was based upon a solved experimental structure (PDB ID: 7NVH)^20^, but because it was missing the entire amphipathic helix, AlphaFold data^47^ was utilized to model this in. Residues 341-377 from the AlphaFold prediction were combined with the cryo-EM structure using PyMOL (Schrödinger, LLC). The root mean squared distribution (RMSD) of the experimentally resolved structure and the prediction was 0.628 Å, showing high similarity in the rest of the structure.

When mutations were performed, they were generated using the mutagenesis wizard on PyMOL. For *M. smegmatis* MmpL3, the following mutations were made for cardiolipin binding site 1 (S1A: T236A/K313A/Y317A/K563A), site 2 (S2A: K375A/T376A/F389A/R677A), site 3 (S3A: W737A/R740A/W741A/R744A), and site 4 (S4A: R392A/K399A/R400A and S4EEE: R392E/K399E/R400E). For *M. tuberculosis* MmpL3, the S4EEE (K387E/K394E/R395E) mutant was generated.

CGMD simulations were performed using the Martini 3 forcefield^48^. The protein was converted to this resolution using martinize2^49^, using an elastic network with a force constant of 500 kJ mol^-1^ nm^-2^, and upper and lower elastic bond cut-offs of 1.0 and 0.5 nm, respectively. The protein was positioned and embedded within the membrane using memembed^50^ and insane^51^, generating new starting positions for each simulation. The membrane had a composition of PE:PG:CL = 75:20:5 on both leaflets. The box size (x, y and z) for these systems was ∼12 x 15 x 16 nm. The systems were solvated with Martini water and neutralized using NaCl to a concentration of 0.15 M, resulting in system sizes of ∼25,000 particles. Systems were then energy minimized using the steepest descent algorithm.

A timestep of 20 fs was used for all production simulations in an NpT ensemble at 310 K. The Parrinello-Rahman algorithm^52^ was used for pressure coupling with a τ_p_ = 12.0 ps in a semi-isotropic manner. The Velocity rescaling algorithm^53^ was used for temperature coupling with protein, lipids and solvent & ions separately. The temperature coupling constant was set at 1 ps. All simulations were performed using GROMACS 2021.4^54^.

For simulations of the *M. smegmatis* and *M. tuberculosis* WT MmpL3, as well as the *M. smegmatis* S4EEE variant, five repeats of 10 µs were performed. For simulations of *M. smegmatis* alanine mutants, and the *M. tuberculosis* S4EEE variant, five repeats of 5 µs were performed. Overall, there is a combined total of 300 µs of simulation time.

Analysis of the lipid binding sites was performed using PyLipID^55^ using the headgroup from CL to detect binding sites. From the WT simulations, high occupancy binding sites containing residues known for CL binding^56^ were chosen to further investigation with mutations in silico and in vitro. For visualization of these binding sites, CG2AT2^57^ was used to convert bound poses in CG to atomistic resolution.

### Determination of intracellular pH using the BCECF-AM dye

To investigate the impact of MmpL3^S4EEE^ on ΔpH in *M. smegmatis* cells, we assessed cellular pH changes using BCECF, a pH-sensitive fluorescent dye activated intracellularly via esterase-mediated hydrolysis of BCECF-AM^58^. Briefly, WT *M. smegmatis* cells with the plasmid pJEB402-*hygR-p_ace_-mmpL3^S4EEE^* integrated at the *attB* site were cultured in LB media until reaching log phase (OD_600_∼1.0), followed by induction of MmpL3 expression using 2% acetamide for 6 hours. Subsequently, cells were pelleted, washed, and resuspended in suspension buffer (25 mM MES, 25 mM MOPS, 100 mM KCl, 90 mM glucose, 0.5 µM BCECF-AM) to a final OD_600_ of 0.3. External pH was adjusted to pH 6.8 by adding appropriate amounts of HCl. Intracellular BCECF fluorescence intensities were promptly measured at 525 nm emission (excitation at 488 nm and 440 nm) using a SpectraMax plate reader. Following a 60-minute measurement period, CCCP was introduced to disrupt the *pmf*, and BCECF fluorescence (I_488_ and I_440_) was monitored for 20 minutes. Intracellular pH values for samples were calculated based on a standard curve of pH-dependent fluorescence ratios (I_488_/I_440_), generated with CCCP-treated samples where internal pH equals external pH.

## Acknowledgements

This project was funded by the Singapore Ministry of Education Academic Research Fund Tier 2 grant MOE2019-T2-1-128 (S.-S.C.). R.H.S. was supported by the NUS Integrative Science and Engineering Program-SCELSE scholarship. C.M.B. was supported by an MRC studentship (MR/N014294/1). This project made use of time on ARCHER2 granted via the UK High-End Computing Consortium for Biomolecular Simulation, HECBioSim (http://www.hecbiosim.ac.uk), supported by EPSRC (grant no. EP/R029407/1). P.J.S. would like to thank the SCRTP at Warwick for use of the computing infrastructure. P.J.S. acknowledges Sulis at HPC Midlands+, which was funded by the EPSRC on grant EP/T022108/1.

## Notes

### Competing Interest Statement

The authors have declared no competing interest.

