## Supporting Information for "Cardiolipin modulation of MmpL3 in mycobacteria"

**Supplementary Figures**

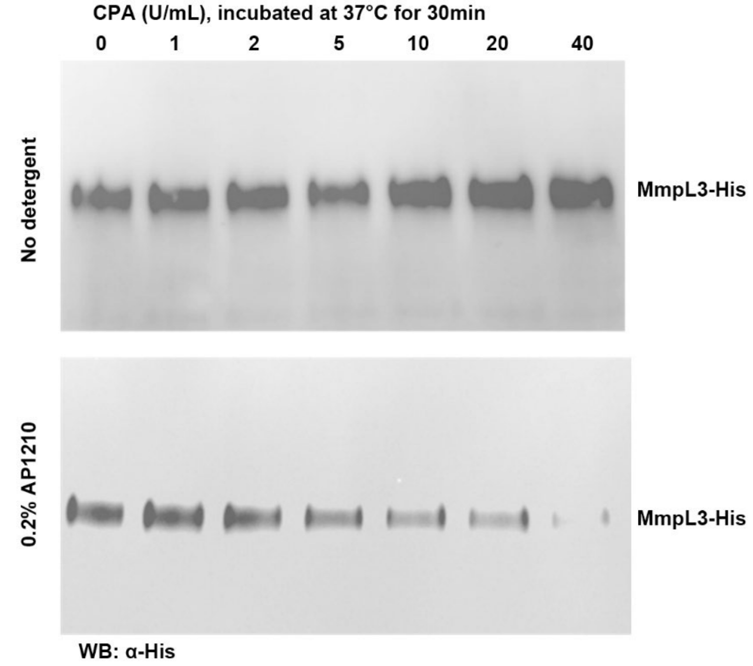


**Figure S1. Purified MmpL3 is reconstituted in the right-side-out manner in proteoliposomes.** α-His immunoblot analyses of purified MmpL3-His reconstituted into liposomes composed of synthetic lipids POPE:POPG (3:1), subjected to C-terminal degradation (loss of cytoplasmic His-tag) by external carboxypeptidase A (CPA) at indicated concentrations. Proteoliposomes were pretreated/solubilized with detergent (0.2% AP1210) (*bottom*) or not (*top*), prior to CPA digestion.

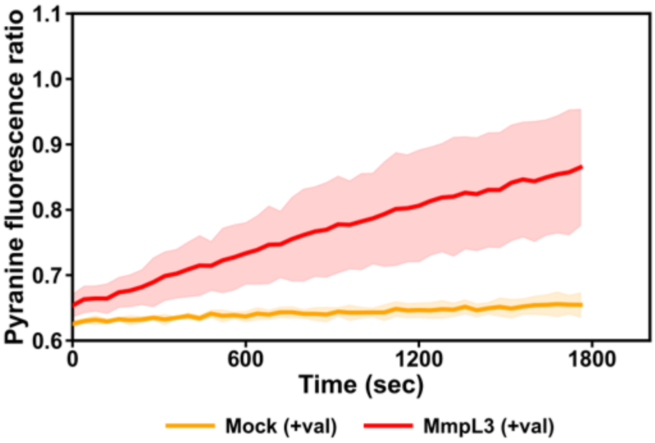


**Figure S2. Purified MmpL3 exhibits proton translocation activity in proteoliposomes comprising synthetic lipids (POPE:POPG = 3:1).** Representative proton translocation assays showing change of pyranine fluorescence ratios (I450/I400) over time of empty (mock) and MmpL3-embedded liposomes due to lumenal alkalization in the presence of an artificial pH gradient (initial $pH_i = 7.0$; $pH_o = 8.0$). MmpL3-His (0.6 pmol) was reconstituted into liposomes composed of synthetic lipids (POPE:POPG = 3:1). Assays were performed in the presence of 1 µM valinomycin as indicated. Means and standard deviations (±SD) of fluorescence ratios from measurements of technical repeats are plotted.

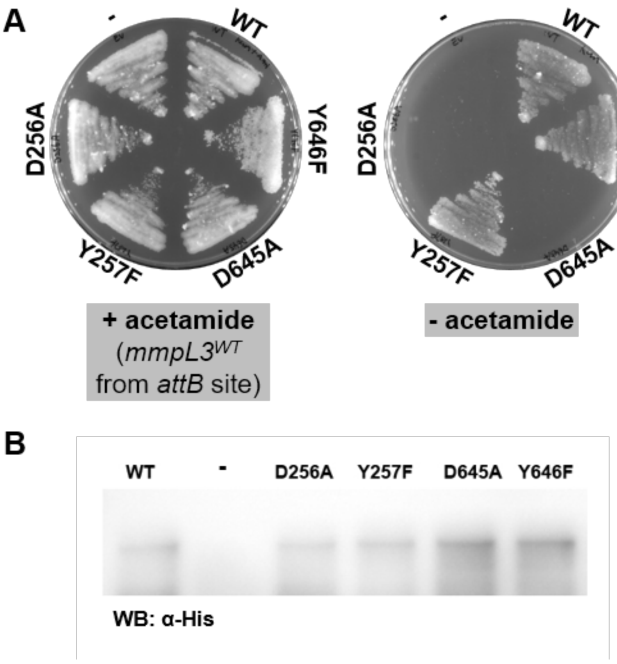


**Figure S3. MmpL3$^{D256A}$ and MmpL3$^{D645A}$ channel variants are non-functional and unable to support growth.** (A) Growth complementation by *M. smegmatis* MmpL3$^{WT}$ or MmpL3 variants (with mutations in channel DY pairs) expressed *in trans* from pMV261 in the *mmpL3* conditional knockout (cKO) strain. Expression of *mmpL3$^{WT}$* from the *attB* site was induced by addition of acetamide (0.2% w/v) where indicated. (B) α-His immunoblot analysis revealing comparable expression of indicated variants in strains in (A) in the absence of acetamide.

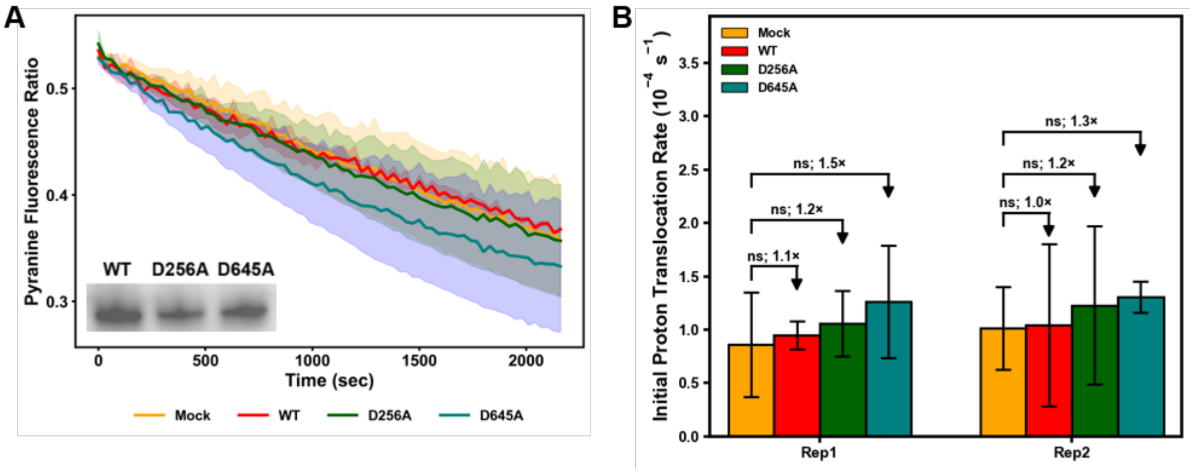


**Figure S4. The proton translocation activities of purified MmpL3$^{D256A}$ and MmpL3$^{D645A}$ channel variants are inhibited by the presence of cardiolipin, similar to MmpL3$^{WT}$.** (A) Representative proton translocation assays showing change of pyranine fluorescence ratios (I450/I400) over time of empty (mock) liposomes and proteoliposomes reconstituted with indicated MmpL3 channel variants, due to lumenal acidification in the presence of an artificial pH gradient (initial $pH_i = 7.0$; $pH_o = 6.0$). Purified MmpL3-His variants (0.6 pmol) were reconstituted into liposomes composed of 20% w/w CL in POPE:POPG (3:1). Inset shows α-His immunoblot analysis of reconstituted MmpL3 proteoliposomes used. Assays were performed in the presence of 1 μM valinomycin. Means and standard deviations (±SD) of fluorescence ratios from measurements of technical repeats are plotted. (B) Initial mean "proton translocation" rates in empty (mock) liposomes and proteoliposomes reconstituted with indicated MmpL3 variants were quantified (from technical repeats such as in (A)) and displayed for 2 of 3 biological replicates (i.e. independently reconstituted proteoliposomes) performed. For each biological replicate, initial rates were calculated by analyzing the initial linear slopes of fluorescence ratio curves, averaged across 2-3 technical repeats. Error bars represent ±SD of initial slopes. Fold changes with or without MmpL3 variant within each replicate are displayed. Student's t-test: ns, not significant.

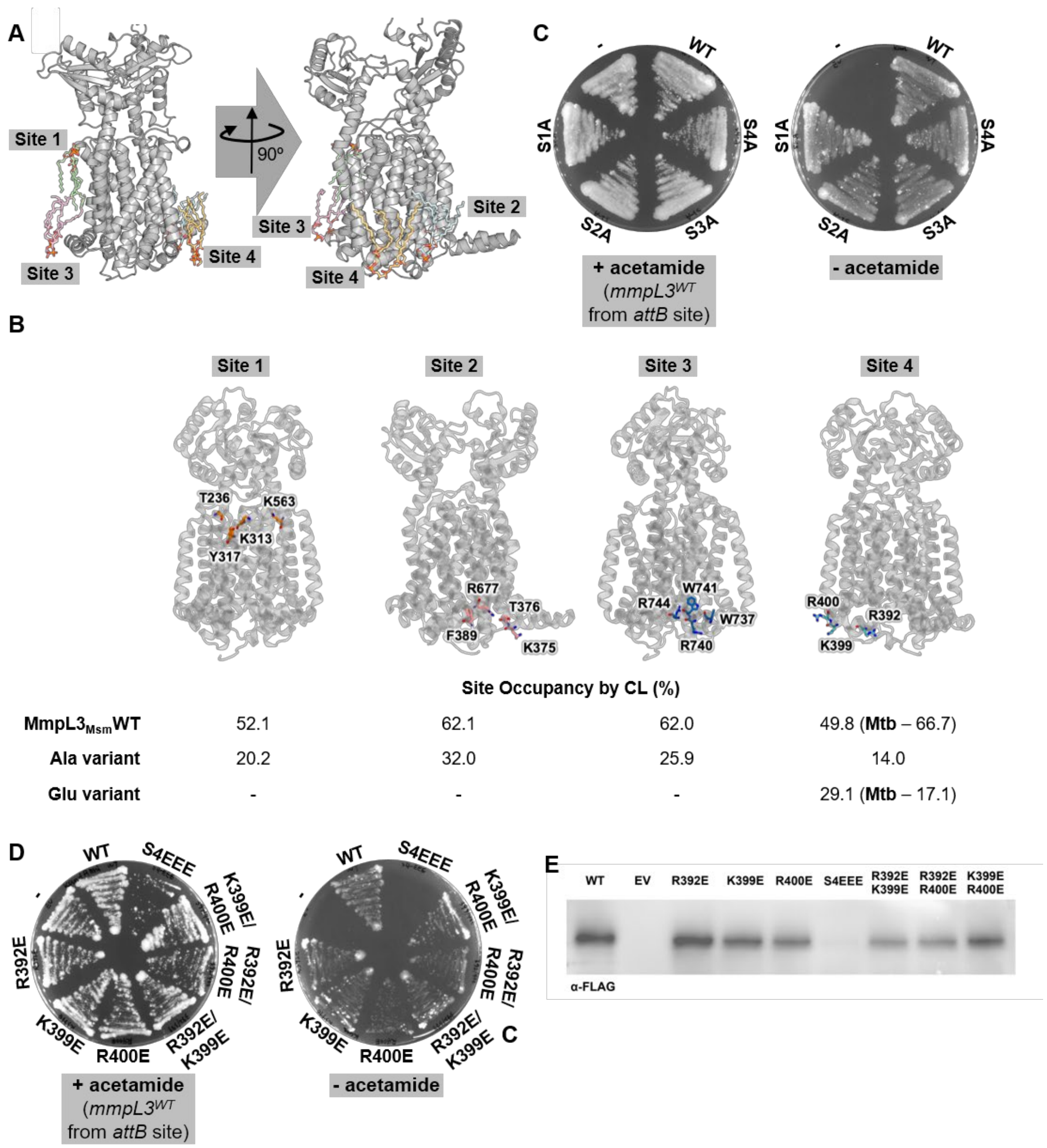


**Figure S5. Charge-swap substitutions in a putative cardiolipin binding site in *M. smegmatis* MmpL3 abolish function.** (A) Cartoon representations of MmpL3 (PDB 7N6B) illustrating putative CL binding sites (sites 1-4) from coarse-grained MD simulations. (B) Specific residues interacting with CL at each site. Site 1 – T236/K313/Y317/K563; Site 2 – K375/T376/R677; Site 3 – W737/R740/W741/R744; Site 4 – R392/K399/R400. In silico CL-occupancy for each site for *M. smegmatis* WT, alanine, and glutamate variants, including indicated *M. tuberculosis* variants, are listed below. (C-D) Growth complementation by *M.*

*smegmatis* MmpL3$^{WT}$ or MmpL3 variants (with (C) alanine or (D) charge-swap substitutions in putative CL binding sites) expressed *in trans* from pMV261 in the *mmpL3* conditional knockout (cKO) strain. Expression of *mmpL3*$^{WT}$ from the *attB* site was induced by addition of acetamide (0.2% w/v) where indicated. MmpL3 variants tested: S1A – T236A/K313A/Y317A/K563A; S2A – K375A/T376A/R677A; S3A – W737A/R740A/W741A/R744A; S4A – R392A/K399A/R400A; S4EEE – R392E/K399E/R400E. (E) α-FLAG immunoblot analysis showing expression levels of indicated variants in strains in (D) in the absence of acetamide.

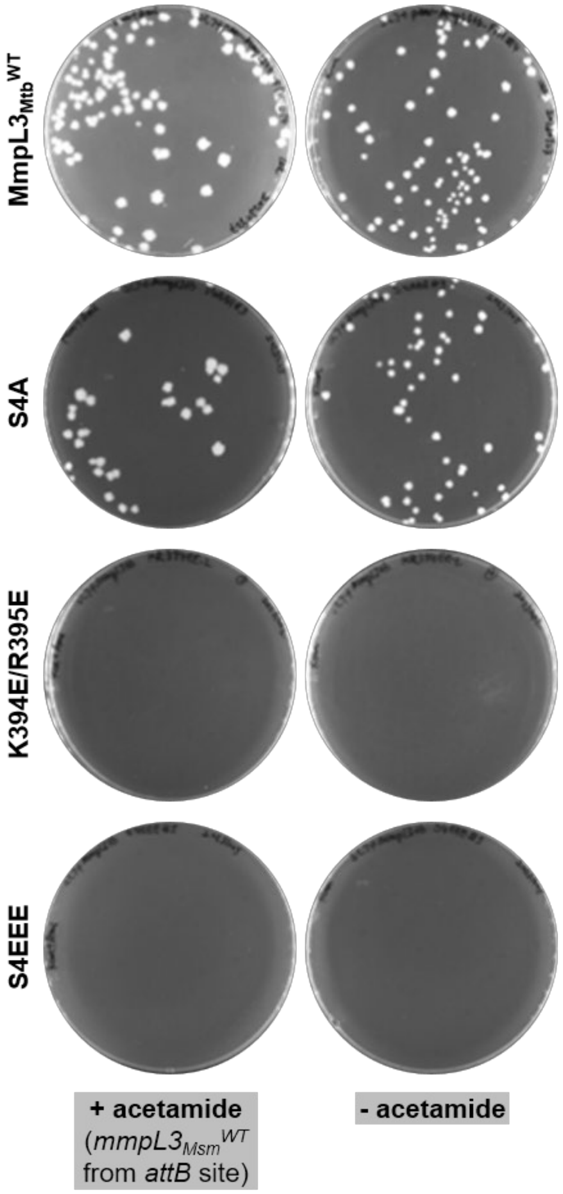


**Figure S6. Charge-swap substitutions in a putative cardiolipin binding site in *M. tuberculosis* MmpL3 abolish function.** Growth complementation by *M. tuberculosis* MmpL3$^{WT}$ or MmpL3 variants (with alanine or charge-swap substitutions in putative CL binding site 4) expressed *in trans* from pMV261 in the *M. smegmatis mmpL3* conditional knockout (cKO) strain. Expression of *M. smegmatis mmpL3$^{WT}$* from the *attB* site was induced by addition of acetamide (0.2% w/v) where indicated. MmpL3$_{Mtb}$ variants tested: S4A – K387A/K394A/R395A; S4EEE – K387E/K394E/R395E.

### Supplementary Tables

**Table S1. MmpL3 occupancy calculations for potential inhibitors in proteoliposomes**

| Potential inhibitors | Inhibitor concentration (μM), $[I]_0$ | Dissociation constant $K_D$ (μM)[a] | MmpL3 concentration (nM), $[P]_0$ | Estimated MmpL3 occupancy[b] |
|---|---|---|---|---|
| AU1235 | 0.1 | 0.29 | 1.175 | ~26% |
| NITD-304 | 0.05 | assumed 0.16 | 1.175 | ~24% |

[a]$K_D$ for AU1235 was determined to be 0.29 μM using microscale thermophoresis[1]. $K_D$ for NITD-304 is not available, but the $K_D$ for related inhibitors ICA-38 or NITD-349 were determined to be 0.16[1] or 0.05 μM[2], respectively. $K_D$ for BM212 is also not available. In another study, $K_D$'s for AU1235 and BM212 were estimated using surface plasmon resonance, and were found to be in the high μM to low mM range[3] – such low affinities are unlikely given that the MICs of these compounds are in the low or sub-μM range.

[b] MmpL3 occupancy is calculated for a 1:1 binding model, where

$$\text{MmpL3 (P)} + \text{Inhibitor (I)} \rightleftharpoons \text{MmpL3-Inhibitor complex (P.I)}$$

Therefore, $[P.I] = \frac{1}{2}\{([P]_0 + [I]_0 + K_D) - \sqrt{([P]_0 + [I]_0 + K_D)^2 - 4[P]_0[I]_0}$

$$\%\text{Occupancy MmpL3} = [P.I]/[P]_0 \times 100\%$$

The estimation here assumes 3D diffusion, but inhibitors are very likely to partition into the bilayer, so true MmpL3 occupancy is expected to be much higher, due to much higher concentrations in the membrane.

**Table S2. List of plasmids used in this study**

| Plasmid | Description | Source |
|---|---|---|
| pJEB402-*kanR-MOP-mmpL3* | integrative plasmid, constitutively express $MmpL3_{Msm}$ in *M. smegmatis* from *attB* site via *MOP* promoter; kanamycin resistance | 4 |
| pJEB402-*hygR-*$p_{ace}$*-mmpL3* | integrative plasmid, conditionally express $MmpL3_{Msm}$ in *M. smegmatis* from *attB* site via acetamide-inducible promoter; hygromycin resistance | this work |
| pMV261-*kanR* | episomal plasmid with *hsp60* promoter; kanamycin resistance | lab collection |
| pMV261-*kanR-hsp60-mmpL3-FTV* | episomal plasmid constitutively expressing *mmpL3-FLAG-TEV-His* in *M. smegmatis* | this work |
| pET22/42-*mmpL3-His* | plasmid for overexpression of *M. smegmatis mmpL3-His* in *E. coli* (λDE3); ampicillin resistance | 4 |

**Table S3. List of primers used in this study**

| Primer | Sequence (5'→3') |
|---|---|
| MmpL3 D256A_fwd | ATCGCGATCGCCTACGGCCTGTTCATC |
| MmpL3 D256A_rev | GGCCGTAGGCGATCGCGATACC |
| MmpL3 Y257F_fwd | GATCGACTTCGGCCTGTTCATCGTG |
| MmpL3 Y257F_rev | ACAGGCCGAAGTCGATCGCGATACC |
| MmpL3 D645A_fwd | TGTCGACCGCCTACGAGGTGTTCCT |
| MmpL3 D645A_rev | CCTCGTAGGCGGTCGACAGACC |
| MmpL3 Y646F_fwd | GACCGACTTCGAGGTGTTCCTGGTG |
| MmpL3 Y646F_rev | ACACCTCGAAGTCGGTCGACAGACC |
| MmpL3 R392E_fwd | TCTGGGGCGAGCTGGTCAACGTC |
| MmpL3 R392E_rev | TGACCAGCTCGCCCCAGAAGCC |
| MmpL3 K399E_fwd | CGTCGTGATGGAGCGCCCGATCGC |
| MmpL3 K399E_rev | CGGGCGCTCCATCACGACGTTGACC |
| MmpL3 R400E_fwd | TGATGAAGGAGCCGATCGCGTTC |
| MmpL3 R400E_rev | ACGCGATCGGCTCCTTCATCACGAC |
| MmpL3 K399E/R400E_fwd | GTCGTGATGGAGGAACCGATCGCGTTC |
| MmpL3 K399E/R400E_rev | ACGCGATCGGTTCCTCCATCACGACGTTGACC |
